# Dodecagon light-sheet fluorescence microscopy for large-volume imaging without striping artifacts

**DOI:** 10.64898/2026.06.29.735400

**Authors:** Chia-Ming Lee, Po-Yen Lin, Xuejiao Tian, Yijuang Chern, Chih-Jen Cheng, Bi-Chang Chen

**Affiliations:** Research Center for Applied Sciences, Academia Sinica, Taipei 11529, Taiwan; Institute of Cellular and Organismic Biology, Academia Sinica, Taipei 11529, Taiwan; Nanoscience and Technology Program, Taiwan International Graduate Program, Academia Sinica 11529, Taiwan; Department of Engineering and System Science, National Tsing Hua University, Hsinchu 300, Taiwan; Institute of Biomedical Sciences, Academia Sinica, Taipei 11529, Taiwan; Division of Nephrology, Department of Internal Medicine, Carver College of Medicine, University of Iowa, Iowa City, IA, U.S.A.

**Author notes:** These authors contributed equally to this work (Chia-Ming Lee, Po-Yen Lin).

**Keywords:** Light-sheet microscopy, Striping artifacts, Tissue clearing, Expansion microscopy

## Abstract

Light-sheet fluorescence microscopy (LSFM) has revolutionized biological imaging by enabling high spatial and temporal resolution with minimal photodamage. However, conventional LSFM techniques often suffer from striping artifacts in the resulting images due to light scattering and absorption within samples, leading to uneven illumination that negatively impacts the accuracy of subsequent image analyses. To address this limitation, we introduce dodecagon light-sheet fluorescence microscopy (dodecaLSFM), a novel approach that maximizes angular diversity to achieve homogeneous illumination and suppress striping artifacts. dodecaLSFM employs diffraction optics and cylindrical lenses to generate twelve light sheets, providing 360° omnidirectional illumination that significantly enhances illumination uniformity compared to traditional mSPIM, mDSLM, and ultramicroscopy systems, which use only one or two illumination planes. We demonstrate the effectiveness of dodecaLSFM by achieving high-resolution imaging of whole mouse brain vasculature following tissue clearing, allowing precise morphometric analysis of vascular networks without striping artifacts. Furthermore, we show that combining dodecaLSFM with expansion microscopy (ExM) enables whole-organ 3D imaging at cellular resolution. This novel approach provides an advanced, scalable solution for large-volume imaging, facilitating detailed structural and functional studies across diverse biological applications.

## 1 Introduction

Imaging cellular interactions and architecture in organisms is an essential aspect of biological research, since it allows cell connectivity and dynamics to be elucidated at the whole-organism scale [1, 2]. Novel developments in imaging methods are vital to deciphering these fundamental processes in biological systems. Light-sheet fluorescence microscopy (LSFM) or selective plane illumination microscopy (SPIM) have become indispensable imaging platforms in many biological fields [3-6]. These methodologies enable intrinsic optical sectioning, exhibit low phototoxicity, and allow imaging at high spatiotemporal resolution [7]. These attributes are achieved by confining illumination to a narrow sheet and then imaging via an orthogonally arranged objective lens. In LSFM, a thin light sheet, generated by a cylindrical lens or by rapidly sweeping a collimated beam, is used to scan the sample plane-by-plane to form a 3D image. When a sample is illuminated by such a light sheet, non-homogeneous background often occurs due to the presence of strongly light scattering or absorbing objects, resulting in striping artifacts [8]. Such non-homogenous illumination significantly degrades image quality and can lead to inaccurate image interpretation [9].

To overcome this issue, several methods have been developed to eliminate striping artifacts. For example, self-reconstructing beams, such as a Bessel beam, can help to reduce scattering artifacts relative to the Gaussian beam conventionally used in LSFM [10]. A Bessel beam can be generated by means of an annular apodization mask [11], an axicon lens[12, 13], or a spatial light modulator (SLM) [14], which helps ensure that light propagation remains invariant over extended distances. When it is partially blocked by an object, the initial intensity profile of the Bessel beam can be reconstructed beyond the obstruction, thereby reducing striping artifacts. However, consequent signal from an out-of-focus plane may increase image background, curtailing accurate image analyses. Imaging of a sample from multiple views represents an alternative approach to suppressing striping artifacts [15,16]. It is achieved by recording a three-dimensional (3D) image stack of the sample from different viewing angles. These multi-view 3D images are then combined using an image fusion algorithm to generate an improved image [17]. However, in multi-view LSFM, the sample often suffers from excessive illumination, which may cause photobleaching. Furthermore, since the sample is usually mounted on a slowly rotating stage and imaged from multiple angles to achieve full sample coverage, temporal resolution is limited. Moreover, LSFM usually generates massive datasets that entail high computational cost [18].

A straightforward way of removing striping artifacts is via multidirectional selective plane illumination of the sample (mSPIM), whereby the light sheet is tilted in the image plane [5, 19] or two parallel light sheets from opposite sides of the sample are used for illumination [20], thereby reducing shadows. In conventional LSFM, the light sheet propagates in roughly the same direction due to the low numerical aperture (NA) configuration. The resulting lack of angular diversity can promote striping artifacts, but this issue can be alleviated by deploying mSPIM. Moreover, mSPIM can be combined with digital scanned light-sheet microscopy (mDSLM) to increase angular diversity in the image plane by utilizing an elliptical Gaussian beam [21]. However, despite this, angular diversity remains limited and striping artifacts can persist for low signal intensity or dense samples even when mSPIM is applied [22].

In this study, we adopted a direct approach to maximizing angular diversity and thereby reducing striping artifacts, termed dodecagon light-sheet fluorescence microscopy (dodecaLSFM), which utilizes twelve light sheets to illuminate samples. Relative to mSPIM, mDSLM or ultramicroscopy approaches, all of which use one or two lenses to generate the illumination plane, our dodecaLSFM method exploits diffraction optics and cylindrical lenses to create twelve light sheet planes, resulting in homogeneous illumination covering a total angular range of 360°. We also demonstrate that combing dodecaLSFM with expansion microscopy represents an effective technique for whole-organ 3D imaging at cellular resolution without striping artifacts.

## 2 Methods

### 2.1 Microscope setup

As illustrated in Figure 1 (Supplement 1, Visualization 1), three laser beams with excitation wavelength of 488 nm, 561 nm and 647 nm are combined and expanded to a diameter of 6 mm using two-lens telescope configuration. Each beam is split into twelve equally intense beams using a diffractive beam splitter (MS-356-561-Y-X, Holo/Or) mounting on a goniometric stage (GNL10, Thorlab). The diffractive optical element (DOE) is a custom 12-fold ring splitter designed for 594 nm. It divides the incident beam into twelve symmetric sub-beams on a conical surface. The wavelength dependence of the diffraction half-angle was measured by projecting the multicolor beam pattern onto a 5 mm grid placed 1.0 m from the DOE (Supplement 1, Figure S1). The measured ring radii and corresponding half-angles confirm the DOE’s designed linear θ–λ scaling centered near 594 nm. For chromatic compensation, the 12-beam ring would otherwise expand from ≈112 mm (488 nm) to ≈138 mm (640 nm) at the sample plane. An electrically tunable lens (ETL2; Optotune EL-16-40-TC-VIS-20D) positioned 125 mm downstream of the DOE adjusts its focal length so that all wavelengths form a 127 mm-diameter illumination ring at the sample plane. The setting of ELT2 are within the ±10 D tuning range of the lens and maintain the beam footprint (< 8 mm radius) inside the 16 mm clear aperture.

**Figure 1.**
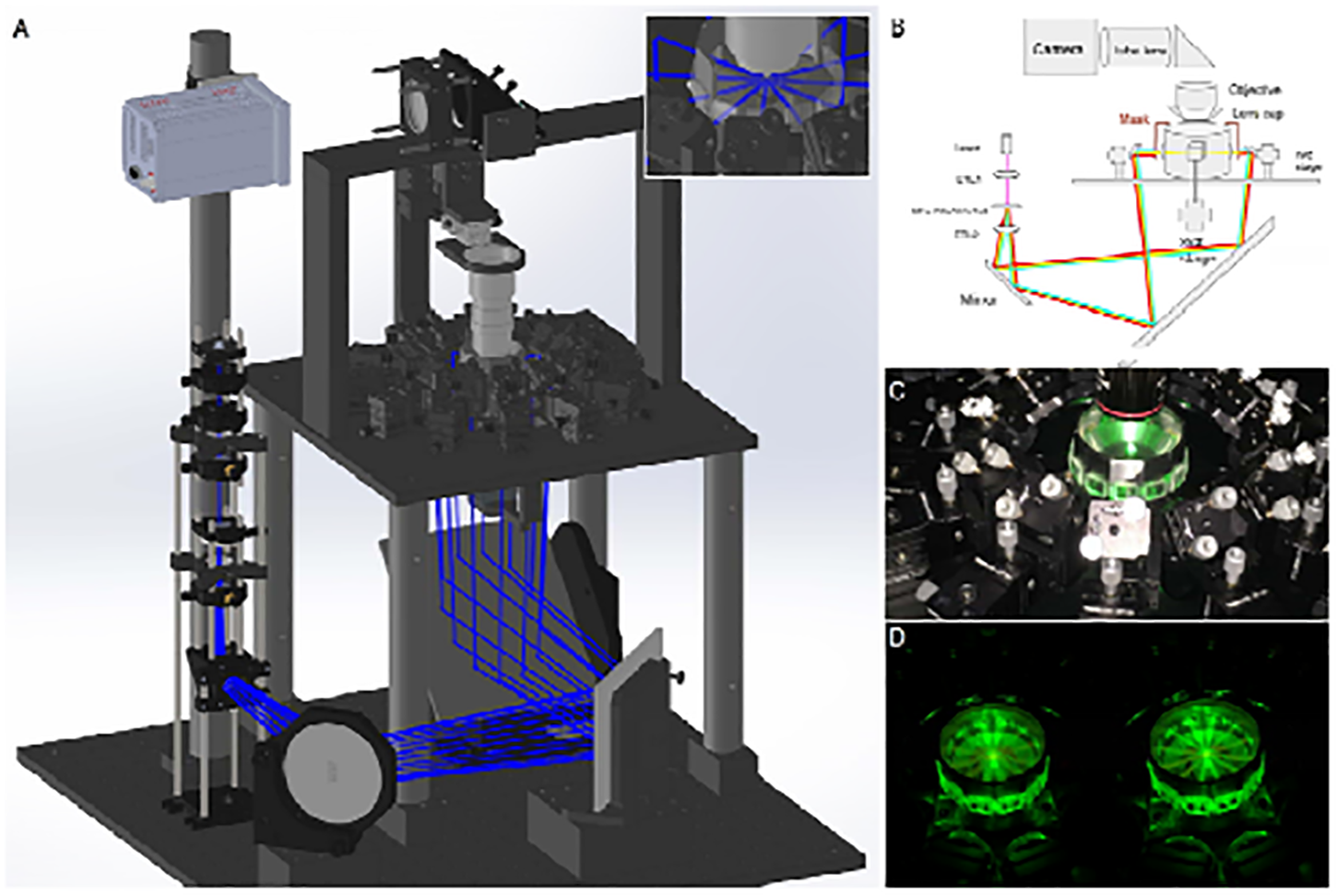
Overall view of the dodecaLSFM setup. (A) 3D rendering and (B) single-plane side-view diagram of the dodecaLSFM model. Laser beams (with paths depicted in blue) emitted from a single optical fiber are split by a custom designed grading plate (MS-356-561-Y-X, Holo/Or Ltd.) to form twelve beams in a circular arrangement. An electrically-tunable lens (ETL2) is used to adjust the angle of the rays, projecting them to an array of mirrors arranged circularly around the sample chamber. (C) Representative photograph of the sample chamber and the surrounding adjustable mirror array. The mask shown in B has been removed for better visualization. (D) Visual representation of the light-sheet illumination for different effective numerical apertures (controlled by slit apertures) to provide different beam widths.

A system of mirrors is used to guide the collimated beams into twelve cylindrical lenses of 100 mm focal length, arranged around the dodecagon-shaped sample chamber, creating twelve light sheets that intersect at the center of the sample chamber. These light sheets can be modulated by adjusting the slit apertures of the lenses to control their thickness and length, rendering them adaptable to varying sample sizes. They are aligned to overlap at their beam waist and positioned at the focal plane of imaging lens. The number aperture (NA) of the light sheets ranges from 0.01 to 0.03. The corresponding light-sheet thickness (at the beam waist) is approximately 10 µm to 30 µm, while the propagation length (approximated by the Rayleigh range) is 500 µm to 4 mm, varying slightly with wavelength and focusing conditions. These parameters were optimized to balance axial resolution and field of view for volumetric imaging. In Table 1, we list the optical parameters and the slit geometry with the resulting resolution estimated by Rayleigh criterion in microns. Custom-made sample holders are designed for various sample types, such as clearing or expansion samples, and mounted at the center of the chamber. For the imaging part, the fluorescent images were detected using standard objective (Olympus, 10X NA 0.6) with tube lens of 200 mm (Thorlabs, TTL 200) or Canon 85 mm Lens with tube lens of 135mm and acquired by the camera (Hamamatsu, V2). The sample is placed in the dodecagonal chamber containing a refractive index (RI)-matching buffer and it is translated using a triple-axis translation stage for large-volume imaging. dodecaLSFM enables multicolor 3D imaging, which is achieved by imaging the same position sequentially with excitation lasers of different wavelengths and using corresponding emission filters (Supplement 1, Figure S2). This capability enables determinations of the relative distributions of various organelles within tissues.

**Table 1.** Axial resolution optical parameters and the slit geometry.

| Slit diameter (mm) | NA | Estimated resolution | Rayleigh range |
| --- | --- | --- | --- |
| 0.5 | 0.01 | 28.1 $\mu\text{m}$ | 4.4 mm |
| 1 | 0.02 | 14.0 $\mu\text{m}$ | 1.1 mm |
| 1.5 | 0.03 | 9.4 $\mu\text{m}$ | 0.5 mm |

Cylinder lens focal length = 25 mm, $\lambda$ = 488 nm

To characterize the resolution of our microscope, we imaged 0.5□µm fluorescent beads embedded in a gel matri using a 1.5 mm slit diameter configuration. Representative lateral and axial point spread functions (PSFs) are show (Supplement 1, Figure S3). Based on the measured PSFs, the lateral and axial resolutions were estimated to be approximately 9 µm and 9 µm, respectively. These values are consistent with expectations given our system’s optical configuration. The results also indicate that the twelve light sheets are well-aligned axially, with no significant degradation in axial resolution due to misalignment

### 2.2 Sample preparation

For CLARITY-cleared whole mouse brain and vascular imaging, DyLight 649-conjugated L. esculentum (Tomato) lectin (LEL-D649, no. DL-1178, Vector Laboratories) was used to label vasculature during tissue clearing process. Adult mice (4-12 weeks old) were deeply anesthetized with a mixture of ketamine/xylazine (100/20mg/kg). LEL-D649 was diluted in saline to a concentration of 0.5 mg/mL and injected into the body of an anesthetized mouse. Mice were then perfused transcardially with ice cold 1X PBS followed by a mixture of 4% (wt) PFA, 4% (wt/vol) acrylamide, 0.05% (wt/vol) N, N’-methylenebisacrylamide, 0.25% (wt/vol) VA-044 and PBS. CLARITY cleared mouse brains were processed as previously described [23]. In brief, the freshly dissected brains were then incubated in the same solution at 4 °C for 3 days, followed by washing with 1X PBS for 3 times, with 30 min each time. Samples were polymerized with the same gelling solution at 37 °C for 2 hours. Lipid extraction was performed in a SDS solution containing 6% SDS and 50 mM Tris with passive thermal diffusion method at 37 °C for 30 days with gentle shaking. SDS solution was refreshed every 5 days, and the mouse brain samples should become nearly transparent in the SDS solution after lipid extraction step. For refractive index matching and imaging, RapiClear 1.49 was used. CLARITY-cleared samples were incubated in a first 10-mL of RapiClear 1.49 solution for 1 day at room temperature with gentle shaking, followed by a second 10-mL RapiClear 1.49 incubation at room temperature for additional two days with gentle shaking. Before imaging, sample was transferred to the imaging chamber with 50-mL fresh RapiClear 1.49, and remained in the chamber for overnight.

For Immunostaining, the fixed samples were first blocked 1 hour at 37°C in blocking buffer containing 10% normal goat serum (NGS) and 0.2% Triton X-100 in 1× PBS. Then, they were incubated with primary antibody (rabbit anti-Nkcc2, 5 µg per 7-day old mouse kidney; AB2281) in immunostaining buffer [1X HEPES-TSC (10mM HEPES, 10% Triton X-100, 1.17% w/v NaCl, 0.5% w/v Casein), 2.5% Quadrol, 0.5M Urea prepared in ddH2O] for 3 days, followed by incubation with secondary antibody (donkey anti-rabbit Alexa Fluor 488, 1:100 dilution; Invitrogen A21206) in immunostaining buffer for an additional 3 days. All antibody incubations were carried out at room temperature with gentle shaking. After each antibody incubation, samples were thoroughly washed with 0.1 M PBT solution [0.1M PB (1.2% w/v Sodium phosphate, 0.24% w/v Sodium dihydrogen phosphate dihydrate, 0.01% Sodium azide) pH 7.5, 10% Triton X-100, 10% Sodium azide prepared in ddH2O] for three times with 1 hour each time. A final wash was performed in 1× PBS for 2 hours following secondary antibody incubation. Samples were then post-fixed in 1% formaldehyde (FA) for overnight at room temperature, followed by three additional washes in 1× PBS for 1 hour each.

For 4X-Potassium acrylate (KA) expansion microscopy, immunostained and 1% FA fixed samples were firstly pre-incubate in 4X-KA-ExM gelling solution for 2 days at 4 °C as previously described [24, 25]. For 4X-KA gel formation, the gelling solution was made fresh by combining the stock monomer solution and 1% of the initiator VA-044. The gelation occurred in a gelation chamber by placing the sample in the middle of two 15-mm coverslips, and 200 µL of the freshly made gelling solution was used for each sample. Samples were placed in a 37 °C hybridization oven for at least 2 hours until the gel completely solidified. Excess gel was trimmed using a scalpel and samples embedded in gel were transferred into SDS denaturation solution (200mM SDS, 20mM Tris) for gentle shaking at 70 °C for 2 days. After SDS denaturation, samples were washed three times in 1X PBST (1X PBS, 0.2% Triton X-100) at 37 °C for 2 hours each time, followed by two times wash in 1X PBS at room temperature for 30 min each time. Finally, samples underwent a water-based expansion (dilation) step for at least 1 day at 37°C with gentle shaking. To maintain structural integrity during imaging, 0.1% low-melting-point agarose in water was used to support the expanded samples, which were mostly water and therefore extremely fragile. To prevent continued expansion, samples were transferred from agarose water to ddH□O at room temperature for at least 2 hours prior to imaging.

### 2.3 Quantification and statistical analysis

To qualify the vasculature in brain and kidney, the tubular structures were first traced using the open source software neuTube [26] and saved into SWC files. The trace results were processed using the centerline tree module in Amira (Thermo Scientific) and the structural characteristics of the vascular network were identified. The vessel length is defined as the segment that connects two branching points. The module then calculated the length based on the path of the centerline between the two connected points. The vessel diameter (or radius) is the estimated cross-sectional width of the structure at a given centerline point, typically calculated based on the surrounding voxel data in the segmented 3D volume. Next, the tortuosity, as defined curved segment length/distance between start and end point of the segment, is a geometric metric that describes how much a vessel deviates from a straight path.

## 3 Results

### 3.1 Suppressing striping artifacts via dodecaLSFM

In order to quantify the impact of light-sheet illumination angular diversity on striping (shadow) artifacts, we analyzed the autofluorescence images of expanded mouse kidney acquired using different light-sheet illumination configurations (Supplement 1, Visualization 2, 3). In Figure 2, we present single-plane images cropped from the whole kidney after 4x expansion, with plots below the images showing the intensity profiles along the red dashed line of the respective images. Following conventional single-plane illumination (Figure 2A), striping artifacts are clearly observable due to sample scattering and absorption, resulting in the reduced signal intensity (∼10%) apparent in the intensity line profile (red arrow). Moreover, striping artifacts persisted when dual light-sheet illumination at an angle of 30°, 90° or 180° was applied (Figure 2B, C, D). The two laser beams of the dual-beam light sheet configuration are usually arranged orthogonally or in opposite directions to reduce striping artifacts, but shadows are still present in the resulting autofluorescence images and they lead to an uneven illumination plane. The consequent reduction in signal intensity is approximately 3 % to 9 % in our images (red arrows). Thus, in these configurations, the lack of illumination angular diversity cannot overcome the challenge of striping artifacts. However, when we imaged the same sample using our dodecagonal illumination array, the intensity reduction at the same position almost disappeared (Figure 2E, red arrow). Thus, dodecaLSFM prevents light scattering and absorption by structures obstructing the laser beam and that can cast a shadow, resulting instead in a uniform illumination plane.

**Figure 2.**
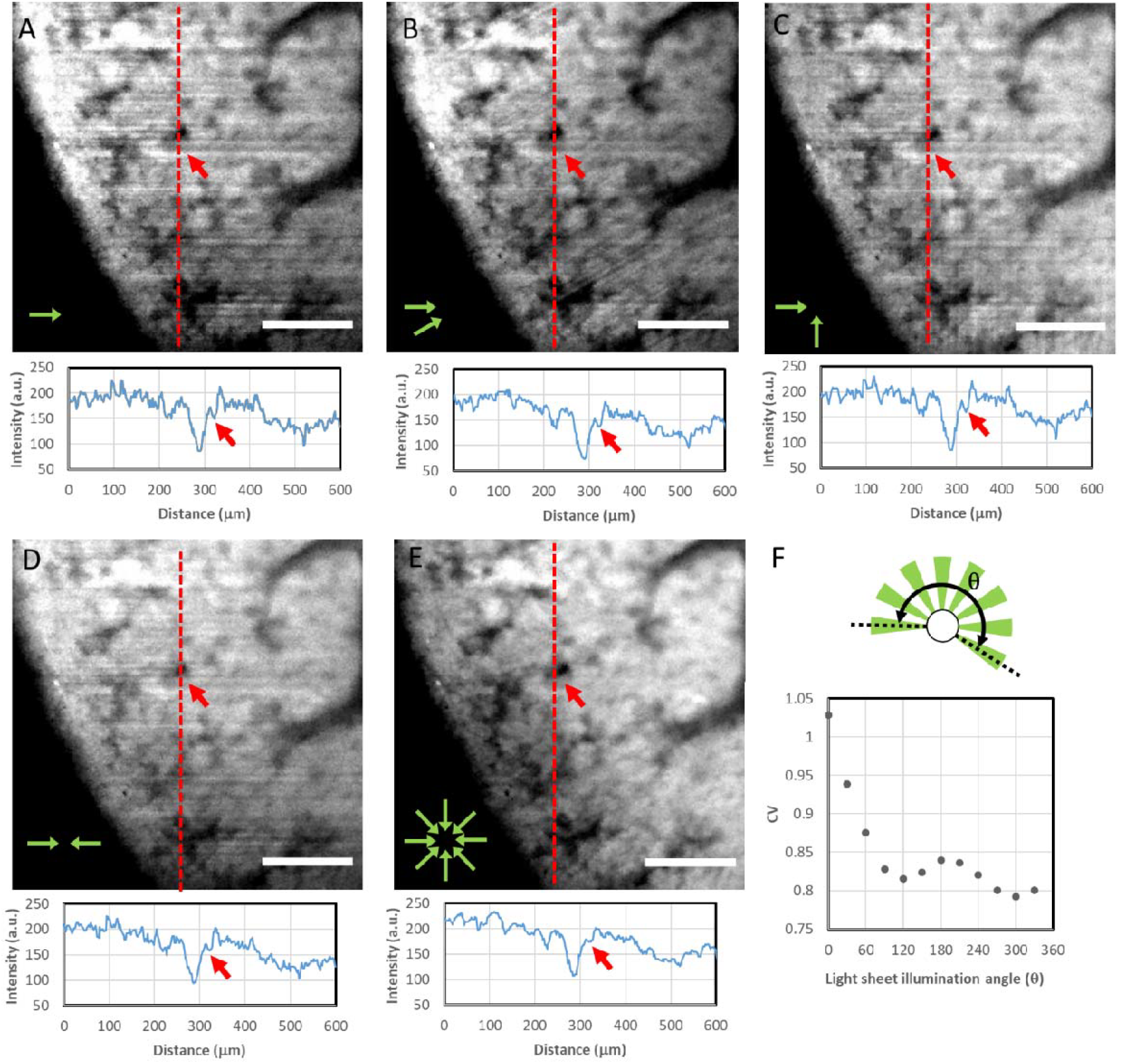
Representative autofluorescence images of an expanded whole mouse kidney acquired by illumination with (A) a single light sheet, (B) dual light sheets at 30°, (C) dual light sheets at 90°, (D) dual light sheets at 180°, and (E) twelve light sheets covering 360°. The plots below the images show the intensity line profiles along the red dashed line. The red arrows indicate the diminished signal intensity that appeared following single or dual light-sheet illumination at different angles that was recovered by dodecagonal illumination. (E) Coefficient of variation (CV) of the signal intensity distribution of autofluorescence images acquired for differing angular variations in light sheet illumination. Scale bars: 200 μm.

Moreover, we determined the coefficient of variation (CV), calculated as the ratio of standard deviation (σ) to the mean (μ) of the image intensity distribution, to verify the impact of the angular diversity of light-sheet illumination on striping artifacts. A high CV indicates high variability or unevenness in the intensity distribution of an image, whereas a low CV indicates more uniform intensity across the image. We detected that CV values declined with increasing angular diversity of light-sheet illumination (Figure 2F). Under single-plane illumination, the CV value is >20 % greater than for dodecagonal illumination (360°), and CV values remained 5 % above dodecagonal illumination even with an illumination angle of 180° (the light-sheet configuration with illumination in opposing directions). We determined that the illumination angle needs to cover at least 270° to remove striping artifacts. These results indicate that greater angular variation of the light sheet reduces irregularities in signal intensity, thereby providing an unbiased illumination plane in light-sheet microscopy and thereby mitigating striping artifacts. While simultaneous illumination from twelve directions offers rapid acquisition and effective stripe suppression in cleared or expanded tissues, it may not be optimal for highly scattering samples. In such cases, sequential multidirectional light sheet illumination could preserve angular independence and enhance optical sectioning. Future improvement of the system may integrate beam switching or laser scanning methods to address this limitation.

### 3.2 Imaging of whole mouse brain vasculature by dodecaLSFM following tissue clearing

To demonstrate the imaging capabilities of dodecaLSFM, we performed high-resolution imaging of the whole mouse brain vasculature following tissue clearing. Tissue clearing ensured tissue transparency and allowed us to deploy 3D imaging techniques to visualize tissue structures in the intact samples. In Figure 3A, we present a representative optical section of Tomato lectin-labeled blood vessels at a depth of 3.6 mm in a whole mouse brain (Supplement 1, Visualization 4). The large-volume brain mosaics were composed by stitching the high-resolution Z-stacks (tiles) that covered the whole brain. In Figure 3B, we show a 3D reconstruction of the region encompassed by a red box in Figure 3A, together with the vessel tracking results generated using neuTube (an open-source neuron tracing software used to extract diameter measurements of vascular structures). If striping artifacts occur in an image they may degrade image quality and prohibit accurate image analyses, such as filament tracking. As illustrated in Figure 3C, the image planes generated by dodecaLSFM at different depths are free of striping artifacts. Thus, our novel imaging approach using tissue clearing and dodecaLSFM improves image quality of the capillary features in whole mouse brain by suppressing striping artifacts, resulting in more consistent and unbiased data.

**Figure 3.**
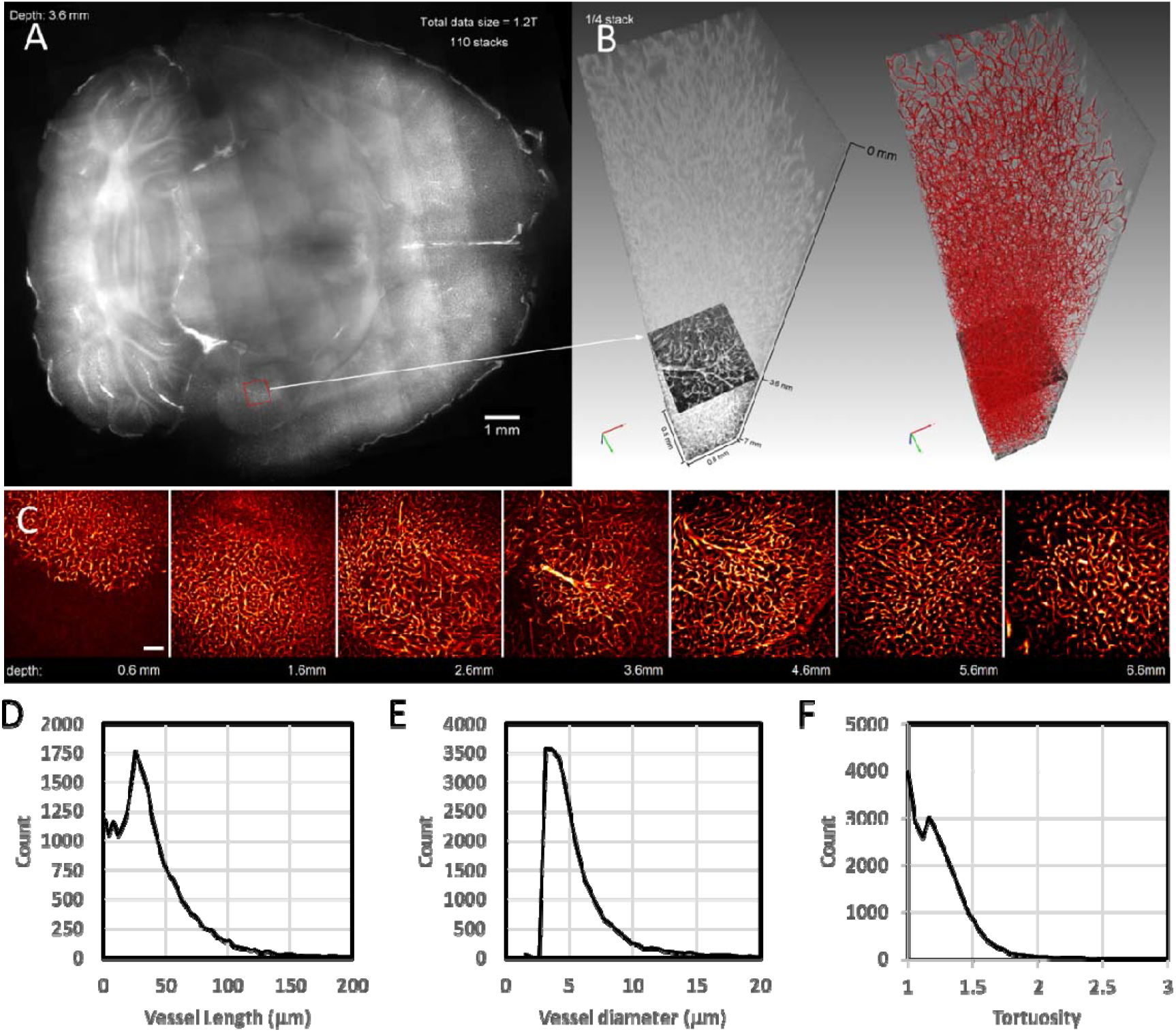
Imaging of the vasculature in whole mouse brain following adoption of the pCLARITY tissue-clearing protocol (see Methods). (A) A representative image of an optical section at a depth of 3.6 mm of DyLight 649-conjugated Tomato lectin-labeled vessels in the whole mouse brain subjected to dodecaLSFM. Scale bar = 1 mm. (B) Magnified volumetric image of the red boxed region shown in (A) and corresponding vessel tracking reconstruction of the vasculature using neuTube. The imaging volume was 800 × 800 × 7000 μm at a voxel size of 0.78 × 0.78 × 5 μm. (C) Images taken at different brain depths (relative to the topmost imaging surface) using a 10× water-immersion objective (Olympus XLPlan N 10× NA 0.6) with a 150 mm tube lens at a voxel size of 0.78 × 0.78 × 5 μm. Scale bar = 100 μm. Quantitative analysis of vessel characteristics: (D) vessel length; (E) mean radius; and (F) tortuosity.

As shown in Figure 3D, we detected a broad distribution of vessel lengths (defined as the expanse between two branching points), with a considerable proportion of short vessels (mean length of 41.2 μm) and a smaller number of long vessels (with lengths reaching up to 200 μm). The distribution of vessel diameters is also biased, with a majority of vessels displaying a diameter of between 3 μm and 10 μm (mean diameter of 6.0 μm), and vessels of larger diameter (>10 μm) accounting for only a small proportion of the total vascular network in the imaged volume. Approximately 85% of the vessels in our brain sample displayed a tortuosity value of between 1.0 and 1.5, indicating that the majority of vessels are relatively straight. These morphometric parameters enable explorations of vascular formation under physiological or patho-physiological conditions and thus our imaging approach can provide the accurate and unbiased measurements of vascular structure essential to diagnosing many diseases.

### 3.3 Imaging of whole mouse kidney with dodecaLSFM expansion microscopy

Next, we applied expansion microscopy combined with dodecaLSFM to characterize kidney development. The resolution of a light microscope is defined as the minimum distance between two or more resolvable features. Expansion microscopy (ExM) is a technique based on physically extending the distance between such features[27, 28], representing a simple method to image features at super-resolution without any need for complex optics. Consequently, ExM combined with light-sheet microscopy has proven an ideal imaging platform for visualizing large volumes at a super-resolution scale. Deploying ExM plus dodecaLSFM, we imaged the Nkcc2-labeled thick ascending limb of the loop of Henle (TAL) in 1-week-old mouse kidneys after 4x expansion and measured the length of the TAL, shown in Figure 4A (Supplement 1, Visualization 5). In Figure 4B, we present representative images at z positions (depths) of 1, 5, and 9 mm, none of which exhibit visible striping artifacts. The tubular structures were analyzed using the Cylinder Correlation module in Amira software, which successfully tracked 3045 individual tubules and provided a comprehensive map of the renal tubular network. Over 90 % of the tubes display a tortuosity value of between 1.0 and 1.5, indicating that the majority of tubules are relatively straight.

**Figure 4.**
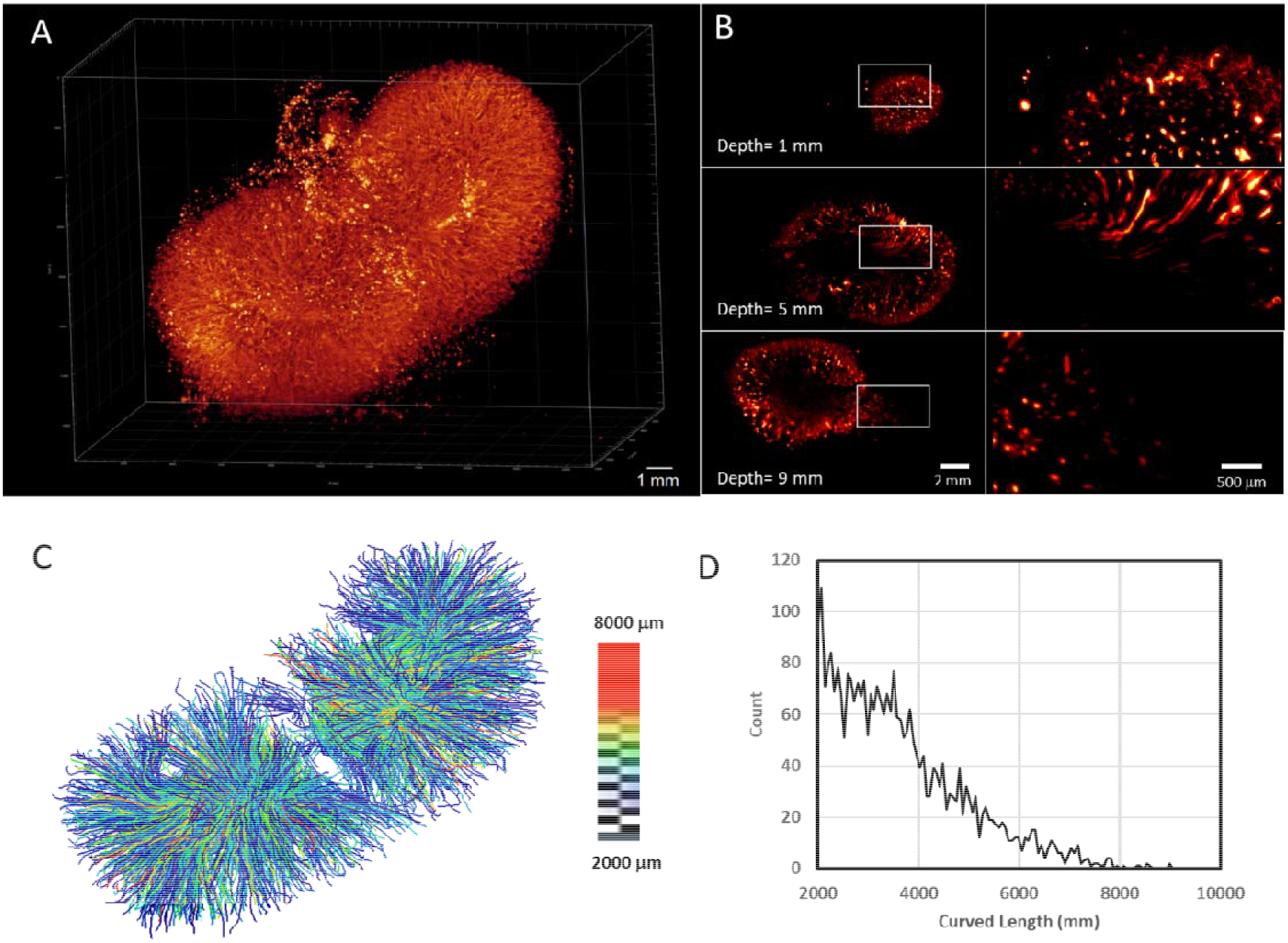
3D dodecaLSFM images from a whole expanded mouse kidney. (A) Immunostaining for NKCC2 in the thick ascending limb of the loop of Henle (TAL) in the kidney. Scale bar = 1 mm. (B) Images captured at different depths (relative to the topmost imaging surface) and magnified views of a cross-section (marked by a white rectangle) using a Canon EF 85 mm 1.4L lens as objective and a Canon EF 75-300 mm as tube lens, at a voxel size of 6.8 × 6.8 × 4 μm (after 4 × 4 binning, imaged using a Hamamatsu ORCA-Lightning sCMOS camera). Scale bars = 2 mm and 500 μm, as shown. (C) 3D reconstruction of the Nkcc2-labeled TAL with color-coded lengths. (D) Length distribution of whole-kidney TALs, ranging from 2 mm to 10 mm.

In Figure 4C, we present a pseudocolored length-encoded 3D reconstruction of the Nkcc2-labeled TAL, enabling determination of the corresponding length distribution across the whole kidney (Figure 4D). Quantitative analysis revealed a broad range of segment lengths, reflecting the complex spatial organization of TAL structures. Expanded sample volumes, particularly those subjected to high expansion ratios, are typically of centimeter scale. To image such large imaging volumes is a unique challenge, as the working distances or depths of field of conventional lenses designed for micron-scale imaging may be insufficient. To accommodate effective imaging of these expanded samples, it is necessary to employ an imaging lens with a long working distance. Our dodecaLSFM system equipped with a macro photographic lens, which is typically characterized by a low numerical aperture, enables imaging of the expanded mouse kidney sample at centimeter scale with single-capillary resolution. With the help of ExM, the imaging capability of our system bridges a critical gap in resolution and field-of-view, allowing us to image large expanded samples without sacrificing the ability to resolve fine structural details. Thus, our combination of a macro photographic lens with dodecagonal light-sheet illumination provides an optimal balance between field-of-view and resolution, with the resulting unbiased illumination proving particularly valuable for studying expanded samples in which both spatial context and microscopic detail are important. Overall, our approach enhances imaging quality and accuracy for large-scale, high-resolution biological analyses.

## 4 DISCUSSION

LSFM is a powerful imaging platform that offers high spatial and temporal resolution while minimizing photobleaching and photodamage. However, traditional LSFM involving single side illumination lacks angular diversity and frequently results in striping artifacts attributable to light-scattering or -absorbing structures within the samples. The consequently uneven illumination can compromise the accuracy of subsequent image analyses. Here, we present dodecaLSFM as a direct approach to maximize the angular diversity of illumination, thereby suppressing striping artifacts. The key advantage of dodecaLSFM lies in its ability to achieve 360° angular illumination coverage, not only improving the homogeneity of light-sheet illumination, but also ensuring more consistent imaging across entire samples. This improvement in illumination uniformity directly leads to higher-quality images, in which sample features can be observed without the interference of striping artifacts. While recent advancements have introduced computational and deep-learning-based methods to digitally remove striping artifacts after image acquisition, relying solely on software-based de-striping for terabyte-scale datasets requires substantial computational resources and prolonged processing times. By fundamentally preventing the formation of shadows through simultaneous 360° optical illumination, dodecaLSFM eliminates striping artifacts during image acquisition, thereby maintaining a simple imaging workflow and minimizing the need for computationally intensive post-processing. This advantage is particularly important for LSFM, which routinely generates terabyte-scale datasets.

Moreover, unlike mSPIM, which relies on sequential multi-angle illumination and image fusion, our approach employs simultaneous illumination from twelve directions to enhance angular diversity in a single acquisition. This simultaneous, stationary acquisition offers distinct mechanical advantages for delicate biological specimens. Traditional multi-view LSFM requires the sample to be mounted on a slowly rotating stage to achieve full angular coverage. This rotation introduces mechanical shear forces that can easily detach, deform, or tear soft tissues. Since both ExM and pCLARITY techniques involve embedding samples in hydrogel matrices that are highly hydrated and extremely fragile, and samples cleared with protocols like CUBIC-R+ also become exceptionally soft, minimizing sample movement is crucial to ensure stable image acquisition. By acquiring 360° illumination simultaneously without rotating the sample, dodecaLSFM minimizes mechanical perturbation and improves ease of use while preserving structural integrity. Although this approach does not fully eliminate asymmetric blur, it enables rapid imaging with effective suppression of striping artifacts in cleared or expanded samples where light scattering is substantially reduced.

Additionally, most commercial light-sheet microscopes that use two or even six static light sheets (e.g., the PhaseView Alpha3 and the Miltenyi Biotec Ultramicroscope Blaze) are excellent platforms for imaging cleared and expanded samples. Their high level of automation enables user-friendly operation for subcellular 3D imaging of large and multiple samples. However, based on our observations, the six illumination sheets in that system do not provide sufficient angular coverage to fully suppress striping artifacts. Our results suggest that an illumination angle coverage of at least 270° is needed to effectively remove such artifacts, particularly in large or optically heterogeneous samples. Apparently, all-directional illumination is the key for larger samples imaging. Moreover, to enable all-directional illumination, the sample mounting system should allow samples to be mounted either by hanging from the top or by being supported from the bottom. In our system, the sample is mounted from the bottom, which provides greater stability, while larger samples may move in both the lateral and axial directions during imaging. The silicone rubber seal between the sample holder and the imaging chamber provides sufficient space for sample translation, while maintaining the imaging buffer within the chamber. However, achieving all-directional illumination still presents challenges in sample accommodation and preparation. While rotation-free imaging provides mechanical advantages for soft pCLARITY and CUBIC-R+ cleared tissues, our observations suggest that CUBIC-R+ buffers may readily crystallize upon exposure to ambient conditions in open-chamber systems. Therefore, careful selection of clearing buffers is required to ensure physical and chemical compatibility with the chamber architecture. Here, we demonstrate an alternative approach to achieving all-directional illumination using 12 light sheets while accommodating the sample effectively.

We have also demonstrated practical applications of dodecaLSFM by performing high-resolution imaging of whole mouse brain vasculature following tissue clearing techniques, as well as of whole mouse kidney following ExM to examine TAL structures in the loop of Henle. Using our protocol, we could visualize vascular structures in the mouse brain, enabling accurate morphometric analyses of blood vessels. Such detailed measurements are vital to assessing vascular formation and structural changes in response to disease [6, 29]. This ability to obtain unbiased measurements of fine-scale features without imaging artifacts is a significant breakthrough in the field of vascular imaging, allowing for more reliable comparisons and analyses of complex biological systems. Additionally, although the use of low-NA optics imposes a limitation on spatial resolution compared to high NA optics, the long working distance and large field of view make them well suited for imaging centimeter-scale cleared or expanded samples. Moreover, the lower NA objectives with long working-distance allows the objectives to be arranged symmetrically around the sample to enable 360° coverage, which is essential for effective stripe suppression in large-volume imaging. While advanced optical configurations, such as lattice light sheet microscopy, axially swept light sheets [30, 31], or the integration of adaptive optics (AO) for real-time aberration correction [32], could further enhance resolution and image quality in deep tissues, they inherently require highly sophisticated, expensive, and maintenance-intensive setups. The aim of dodecaLSFM is to maintain a simple, accessible, and cost-effective system. By relying on a straightforward arrangement of a diffractive optical element and standard cylindrical lenses, dodecaLSFM achieves robust artifact suppression without the high hardware costs and alignment complexity associated with advanced beam shaping or AO. By combining dodecaLSFM with ExM, we demonstrate that 3D imaging of whole organs at a cellular resolution is feasible, making it possible to capture a comprehensive view of tissue architecture in unprecedented detail. This combination of dodecaLSFM and ExM eliminates the problem of striping artifacts that have proven a major impediment to large-scale, high-resolution imaging of biological samples.

## 5 Conclusion

In conclusion, we present a novel and robust dodecaLSFM 3D imaging approach for tissue-cleared and expanded samples at cellular resolution. We developed a dodecaLSFM system to overcome the striping artifacts typically generated by other LSFM methodologies. Its ability to achieve 360° angular coverage provides an even illumination plane and prevents biased image post-processing. It is also compatible with tissue clearing and ExM techniques, offering flexibility in various applications. Although combining with these sample preparation methods can improve the image quality and resolution, these processes are generally not suitable for live imaging, as they involve embedding the sample in a hydrogel matrix and expanding it through physical or enzymatic methods. The resulting improvements in optical transparency and image resolution offer substantial advantages for biomedical research applications. Thus, we believe that the dodecaLSFM technique is a significant step forward for large-scale and high-resolution LSFM imaging with reducing striping artifacts, particularly with respect to studying complex biological structures such as the vascular systems of the brain and kidney. By providing the ability to capture such vascular structures in their entirety at the cellular level, dodecaLSFM opens up new possibilities for exploring both structural and functional aspects of tissues in healthy and disease contexts. This technological approach holds immense potential for advancing our understanding of biological systems, opening up a new frontier in biological research.

## Acknowledgments

B. C. Chen thanks the NSTC and AS for financial support.

## Supplemental document

See Supplement 1 for supporting content.

