## supplemental document 1 for "Dodecagon light-sheet fluorescence microscopy for large-volume imaging without striping artifacts"

**Contents**

**Figure S1.** Compensation of chromatic aberrations in dodecaLSFM.

**Figure S2.** Representative multicolor imaging of an expanded mouse heart using dodecaLSFM.

**Figure S3.** Characterization of the microscope.

**Visualization S1.** Overview of the dodecaLSFM setup.

**Visualization S2.** The dodecaLSFM operated in the multi-angle single-light-sheet illumination mode.

**Visualization S3.** Multi-angle light sheet image of expanded kidney.

**Visualization S4.** 3D visualization of the vasculature in a whole mouse brain cleared according to the pCLARITY protocol, related to Figure 3.

**Visualization S5.** 3D visualization of a whole expanded mouse kidney, related to Figure 4.

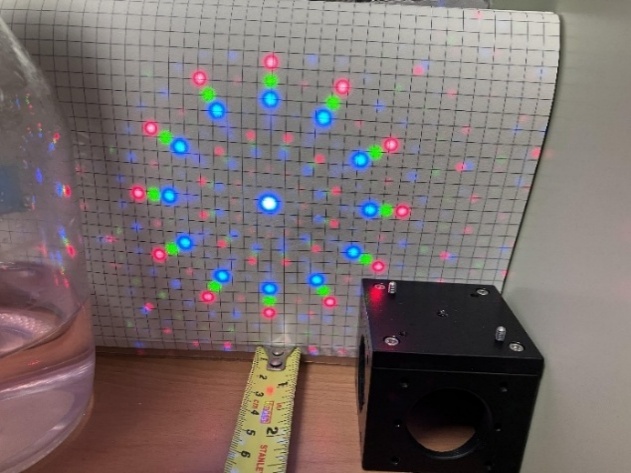

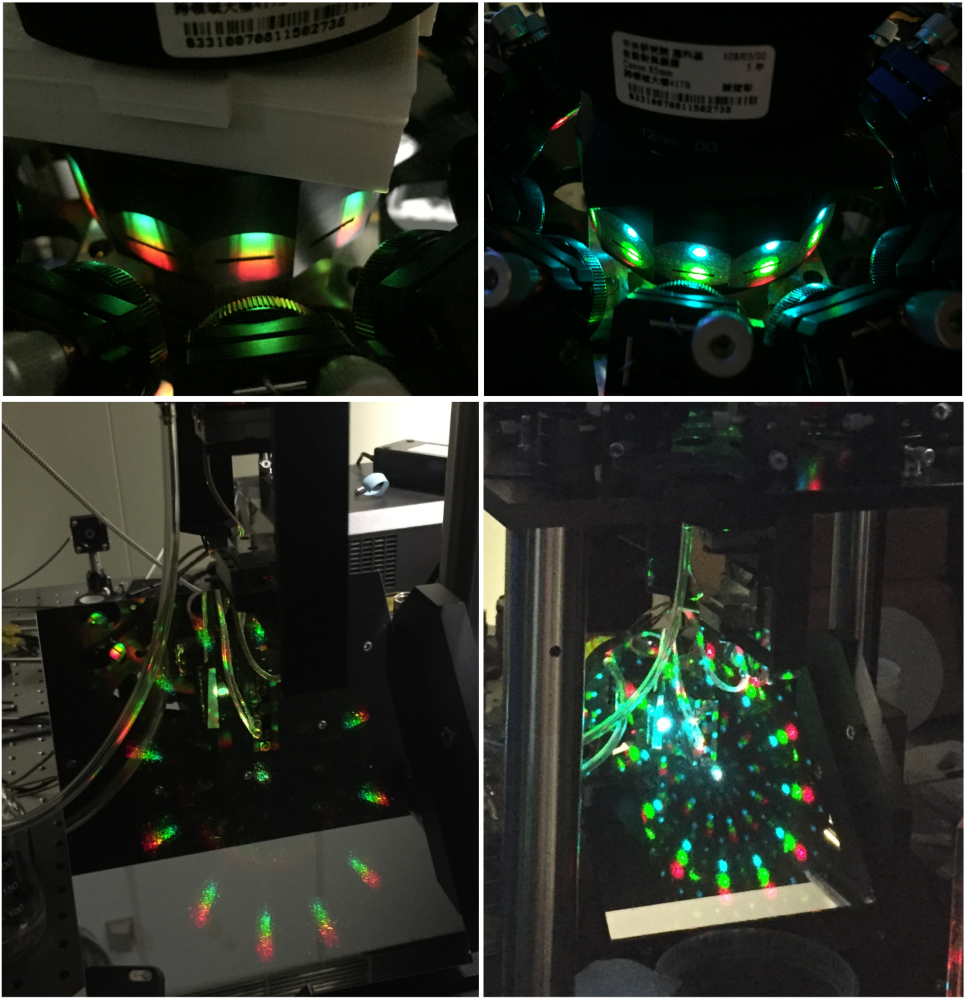

**Figure S1. Compensation of chromatic aberrations in dodecaLSFM**. Left: Experimental projection of the 12-beam DOE pattern onto a 5 mm grid positioned 1 m from the DOE, confirming half-angles of 2.06°, 2.41°, and 2.75° for the 488, 561, and 640 nm wavelength, respectively. Right: Overlay of multicolor beams at the slit aperture after chromatic aberration compensation, demonstrating their spatial alignment.

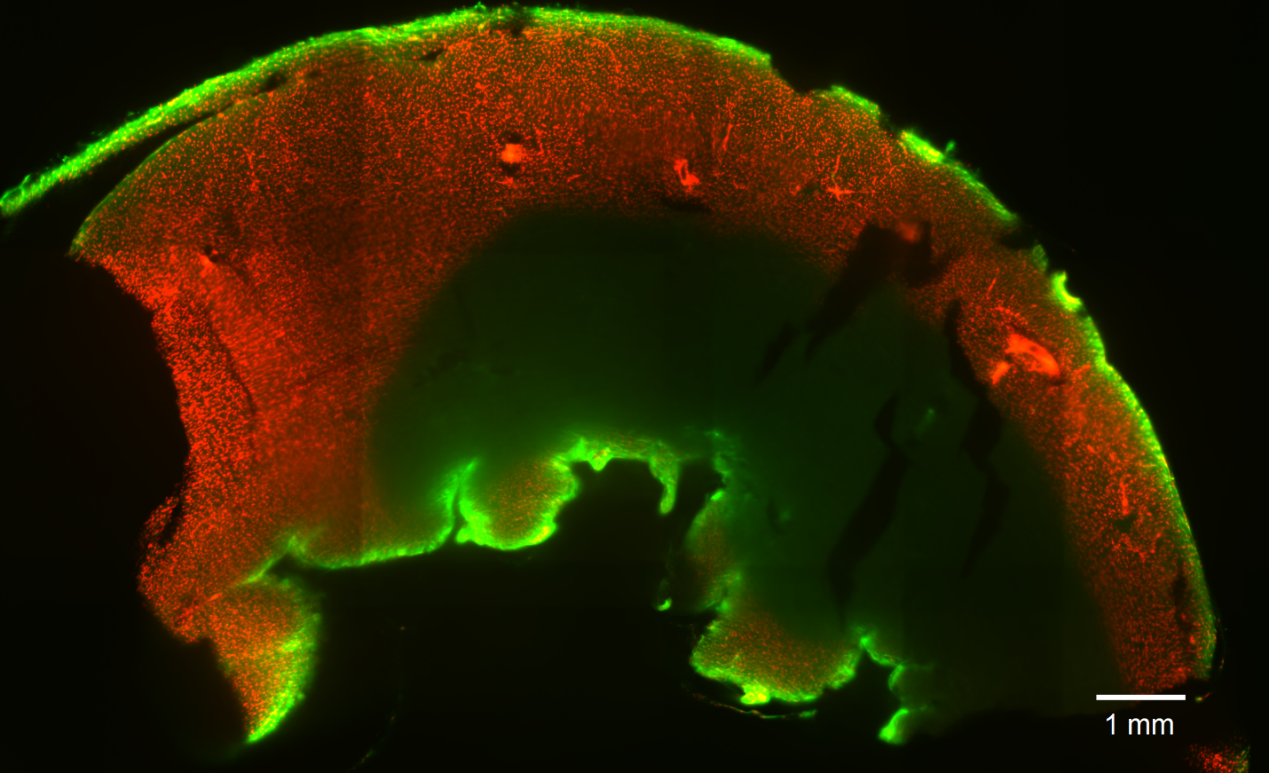

**Figure S2. Representative multicolor imaging of an expanded mouse heart using dodecaLSFM.** The sample was 4× expanded and stained with PI and lectin-649.

.

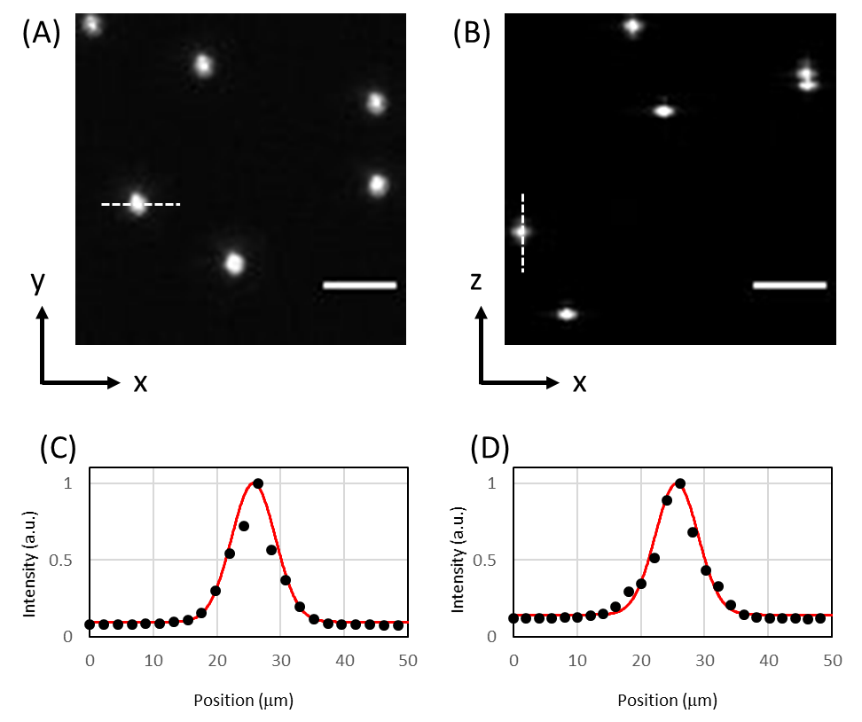

**Figure S3. Characterization of the microscope.** Maximum projections of 0.5 μm bead **image** along XY and XZ are shown in (A) and (B).  The representative Gaussian fit curve of the line profiles of the PSF is shown in (c) and (d); the FWHM is 9.0 μm in X and 9.0 μm in Z, respectively. Black dot: Normalized intensity. Red line: Gaussian fit. Scale bars: 50 μm

**Visualization S1.** Overview of the dodecaLSFM setup.

Visualization S1 present the overview of dodecaLSFM. Laser beam is split into twelve equally intense beams using a diffractive beam splitter and guided into twelve cylindrical lenses of 100 mm focal length, arranged around the dodecagon-shaped sample chamber, creating twelve light sheets that intersect at the center of the sample chamber. The sample is placed in the dodecagonal chamber and it is translated using a triple-axis translation stage for large-volume imaging.

**Visualization S2.** The dodecaLSFM operated in the multi-angle single-light-sheet illumination mode.

Visualization S2 presents the dodecaLSFM operated in multidirectional single-light-sheet illumination mode. The aperture on the rotation stage allows only one excitation beam pass at a time. For 360° omnidirectional illumination, the rotation stage sequentially selects each beam path to illuminate the sample from multiple directions.

**Visualization S3.** Multi-angle light sheet image of expanded kidney.

Visualization S3 demonstrates the stripe artifacts present in autofluorescence images of an expanded kidney in LSFM. Under single light-sheet illumination, pronounced stripe artifacts are observed due to absorption and scattering within the sample.

**Visualization S4.** 3D visualization of the vasculature in a whole mouse brain cleared according to the pCLARITY protocol, related to Figure 3.

Visualization S4 shows the three-dimensional vascular architecture of a whole cleared mouse brain. The reconstructed volume enables visualization and quantitative analysis of the vascular network throughout the entire brain with substantially reduced stripe artifacts, thereby enabling accurate structural analysis.

**Visualization S5.** 3D visualization of a whole expanded mouse kidney, related to Figure 4.

Visualization S5 presents the three-dimensional tubular structures in a whole expanded mouse kidney. 3D reconstruction enables comprehensive tracing and quantification of the TAL network throughout the entire kidney, allowing determination of the segment length distribution. Quantitative analysis reveals a broad range of segment lengths, reflecting the complex spatial organization of the renal tubular system.
